# Nucleation limited assembly and polarized growth of a *de novo*-designed allosterically modulatable protein filament

**DOI:** 10.1101/2024.09.20.613980

**Authors:** Hao Shen, Eric M. Lynch, Chenyang Shi, Joseph L. Watson, Yulai Liu, Kejia Wu, Hua Bai, Chan Johng Kim, William Sheffler, Emmanuel Derivery, James J De Yoreo, Justin Kollman, David Baker

## Abstract

The design of inducibly assembling protein nanomaterials is an outstanding challenge. Here, we describe the computational design of a protein filament formed from a monomeric subunit that binds a peptide ligand. The cryoEM structure of the micron-scale fibers is very close to the computational design model. The ligand acts as a tunable allosteric modulator: while not part of the fiber subunit-subunit interfaces, the assembly of the filament is dependent on ligand addition, with longer peptides having more extensive interaction surfaces with the monomer, promoting more rapid growth. Seeded growth and capping experiments reveal that the filaments grow primarily from one end. We show that designed nucleators that present 12 copies of the peptide ligand promote fiber assembly at concentrations where otherwise assembly occurs very slowly, likely by generating critical local concentrations of monomer in the assembly competent conformation. Following filament assembly, the peptide ligand can be exchanged with free peptide in solution fused to any functional protein of interest, opening the door to a wide variety of tunable engineered materials.

## Introduction

Protein filaments in nature exhibit complex and tunable assembly and disassembly dynamics. For example, microtubules and actin exhibit dynamic instability, where filaments form upon binding of GTP or ATP. Glk1 (Glucose kinase 1) binding to glucose and CTP synthase binding to its substrate (ATP, UTP and glutamine) both allosterically regulate assembly by promoting a filament-competent conformation. IMPDH (Inosine-5′-monophosphate dehydrogenase) binds to regulator adenine and guanine nucleotides directly at an oligomerization interface. PRS1(5-phospho-ribosyl-1(alpha)-pyrophosphate synthetases 1) binds phosphate at its regulator site, which is important for stabilizing the nearby assembly interface (which is completely disordered in the free protomers). Microtubules originate from centrosomes, facilitated by γ-tubulin ring complexes that act as nucleation and anchoring sites (*1*). Actin filaments, on the other hand, nucleate in response to various ligands such as Arp2/3 complex, which binds to existing actin filaments, promoting the formation of new branches (*2*).

Does the complex behavior of natural fibers reflect extremely fine evolutionary optimization, or can it arise in simpler systems? De novo protein design provides an approach to address this question. Computational design methods have been used to generate static fibers from ordered protein monomers, and fibers whose assembly and disassembly are pH dependent (*3*). We reasoned that a designed fiber system in which the monomeric subunit binds an exogenous ligand over an extensive interface independent from the interfaces involved in fiber formation could exhibit ligand-dependent assembly, and provide a simple model system for studying the dynamics of protein filament assembly independent of evolutionary optimization.

## Results

We set out to design self-assembling protein filaments from a designed protein monomer (RPB_PLP3_R6) (*4*) with an extended binding interface for a peptide ligand composed of multiple repeats of the sequence pro-leu-pro (PLP). We chose this system as a starting point because ligand binding affinity can be readily tuned: peptides with 3, 4, 5, and 6 tandem PLP repeats bind with increasing affinity to the monomer, which is composed of six repeating units which each binds one PLP in the peptide. We docked the crystal structure of RPB_PLP3_R6 into protein filament models using the computational method described in Shen *et al.* (*5*), ensuring the peptide binding site was not at any of the filament-generating interfaces (Fig. 1). We generated 91569 helical filament backbones and selected 69 designs for experimental testing based on predicted energy of fiber formation and other metrics (see Methods). The designs were fused to green fluorescent protein (GFP), expressed in *Escherichia coli* under the control of a T7 promoter, and purified by immobilized metal affinity chromatography (IMAC). The peptide ligand (PLP)6 was added to concentrated IMAC eluates and fiber formation was examined by fluorescent microscopy. Most of the designs expressed solubly, and while none formed fibers in the absence of the peptide, one design, HP35, formed fibers upon the addition of the peptide. Negative stain EM examination confirmed the formation of micron-length filaments with the expected morphology (Fig. S1).

**Fig. 1.**
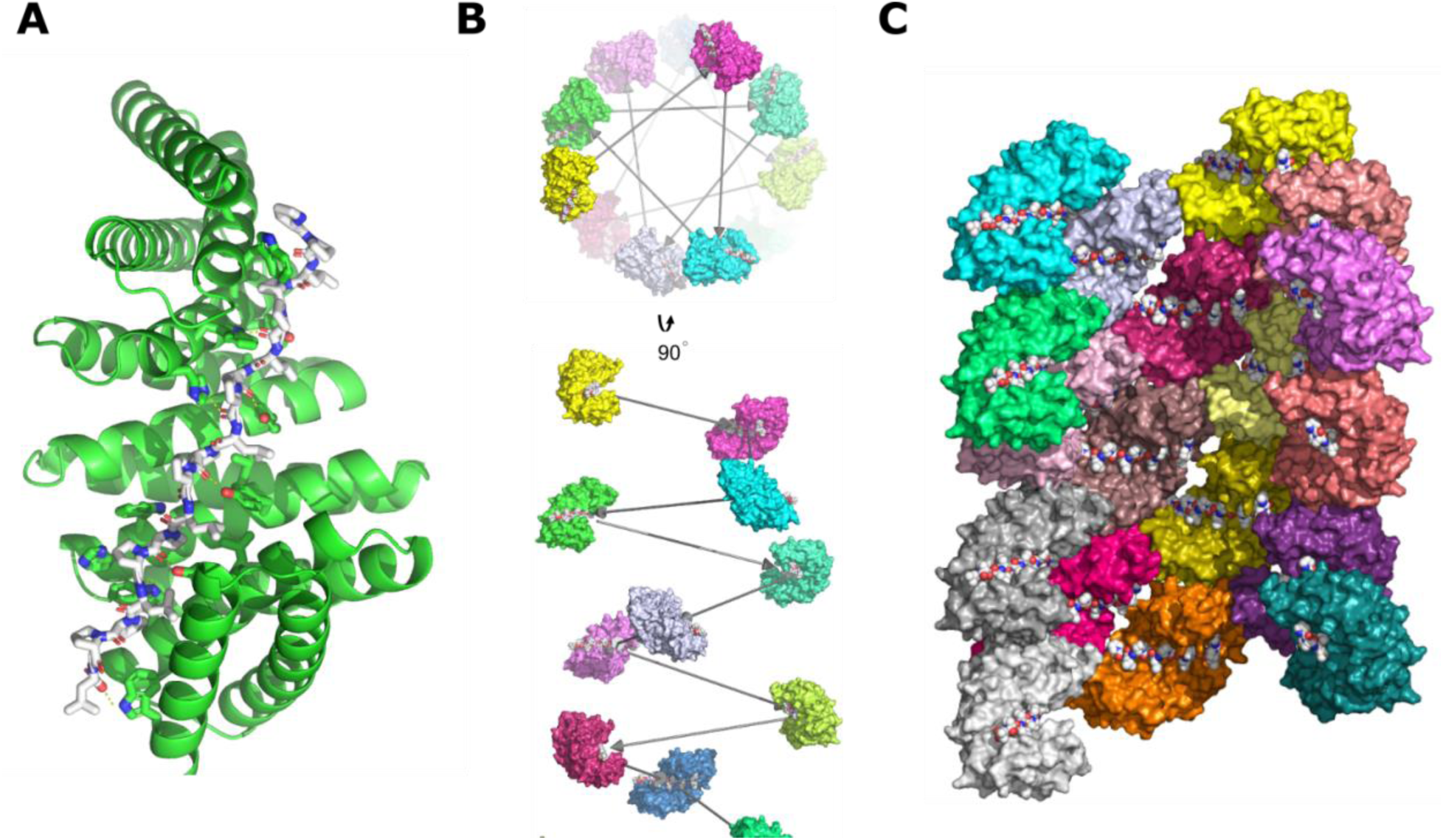
Design strategy for generating peptide-dependent protein filaments. (**A**) Designed peptide binding monomeric starting point (*4*) for fiber design. (**B**) Computational sampling of alternative helical packing arrangements of monomer shown in **A**, with each monomer in a different color. The grey arrows represent the rigid body transform that generates the helical assembly. (**C**) Compact packing arrangement of fibers subunits in lowest energy assemblies following sequence design. Monomers are colored as in **B**.

We froze the HP35-(PLP)6 fibers, generated an initial cryoEM structure using *ab initio* 3D reconstruction in cisTEM (*6*), and performed further helical refinement in Relion (*7*). The structures were solved to 4.0 Å resolution with the protein and the majority of the peptide resolved, with 2.2 Å r.m.s.d. over three chains containing all unique interfaces (Fig. 2). The backbone and side chain conformations at the subunit interfaces are very close to those in the design model. As the peptide is not directly involved in the fiber interface, the peptide dependence on fiber formation must reflect the allosteric effects of the peptide on the conformation of the monomer; the peptide could lock the monomer into an assembly-competent state, or disrupt weak monomer-monomer interactions incompatible with fiber assembly involving the extended peptide binding site.

**Fig. 2.**
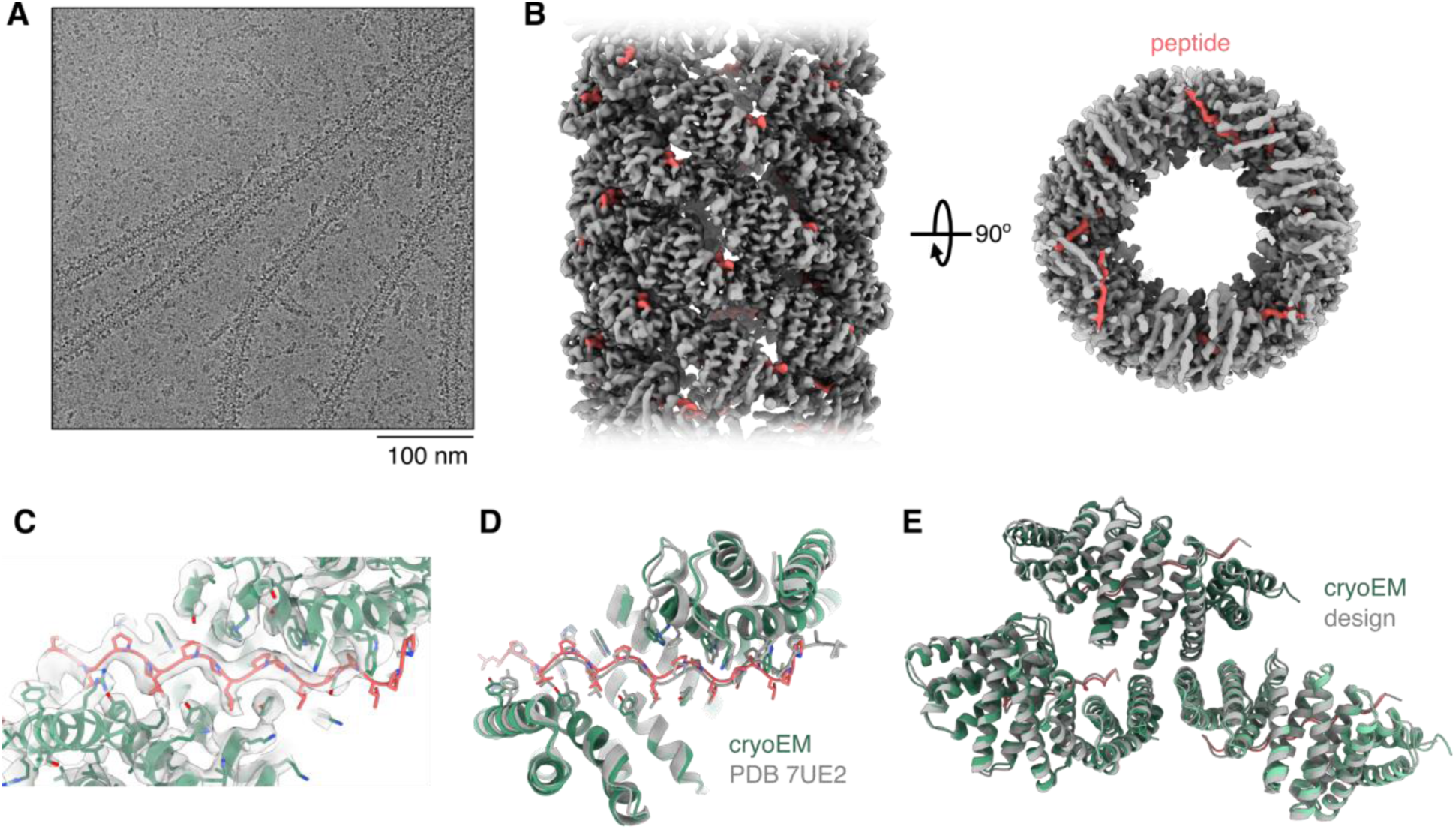
CryoEM structure of HP35. **(A)** Representative cryoEM micrograph of HP35 filaments. **(B)** CryoEM map of HP35. Peptides are colored pink. **(C)** Peptide binding site in the HP35 cryoEM structure. **(D)** Comparison of a monomer from the HP35 filament structure (green and pink) with the original scaffold RPB_PLP3_R6 (PDB 7UE2) (grey). **(E)** Comparison of the HP35 computational design (grey) and cryoEM structure (green and pink) over three monomers comprising the filament interfaces.

We next examined the dependence of fiber formation on the affinity and extent of the monomer-peptide interaction. HP35 fiber formation could be induced with (PLP)4, (PLP)5, and (PLP)6, but not the shorter (PLP)3 (Fig. 3 A and B, Fig S2). The rate of fiber formation increased with peptide length; the half times for 4, 5, and 6 PLP repeat fibers were 1250, 680, and 250 minutes respectively.

**Fig. 3.**
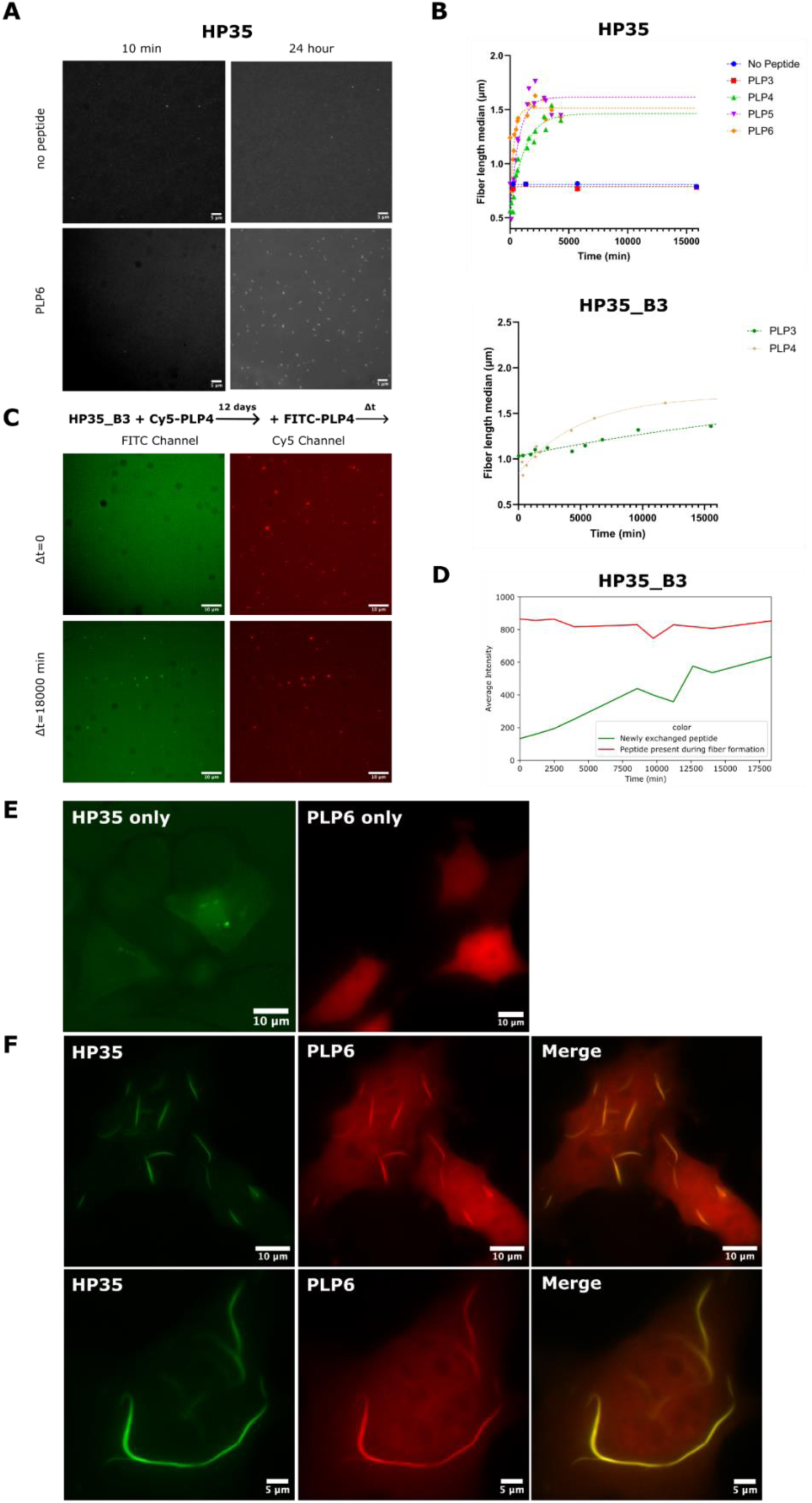
Filament growth dynamics. (**A)** Representative fluorescent microscopy images of HP35 mixed with different lengths of peptide (no peptide vs PLP6) over time and (**B**) quantification of median fiber lengths over time and with the nonlinear fit for HP35 no peptide vs PLP3-6 GFP (top) and HP35_B3 with PLP3-4 GFP (bottom). (**C)** Representative images of HP35_B3 fiber formed by mixing with PLP4_cy5 (top panels) and then PLP4_FITC was added (bottom panels). (**D**) Quantification of PLP4_FITC (green) intensity over time compared to PLP4_cy5 (farred). (**E-F)** Representative images of HP35 fibers in HeLa cells. (**E**) Control images of cells expressing HP35 protein subunit fused to GFP or PLP6 peptide fused to mscarlet-I. (**F**) Representative images of cells expressing both components imaged in green and red channels separately (HP35 and PLP6, respectively) and together (right).

We used a photobleaching experiment to investigate the exchangeability of peptides on fibers. For this experiment, we used a more polar version of the fiber, HP35_B3, with a smaller binding pocket that binds to and forms fibers in the presence of the shorter (PLP)3 and (PLP)4 peptides (Fig. 3B bottom, Fig S3). After forming fiber with cy5 labeled (PLP)4, segments were bleached by laser illumination; after 20 minutes there was no recovery suggesting little or no exchange with unbleached peptide. To monitor peptide incorporation into the fiber over longer times, we added cy5 labeled (PLP)4 to unlabeled HP35_B3 to form the fiber and then added FITC labeled (PLP)4 peptide to monitor incorporation of the new peptide. We found the intensity of FITC increased over hours to days (Fig. 3C and D), indicating that peptides can add to the fiber once formed.

To explore the function of our induced fiber formation system in cells, we expressed HP35 and PLP6 peptide in mammalian cells. Compared with the control cells which only expressed HP35 protein or PLP6 peptide (Fig. 3E), we observed large fibers forming in cells co-expressing HP35 and PLP6 (Fig. 3F).

We studied fiber extension independent of nucleation using pre-assembled GFP-labeled fiber as seeds and then incubated with mKO2-labeled peptide and monomer. We observed robust seeding of growth, with 45% of green fibers having a newly formed red tip (Fig. 4A and B). Unexpectedly, in almost all cases the growth was just from one end: amongst the 740 fibers we analyzed, 340 had one red tip and only 2 had two red tips. Given that 23% of fiber ends (two ends per fiber) have a red tip, we would predict (if the fibers do not have polarity) that the number of fibers with two tips should be 5.3% (0.23*0.23), but the observed value is clearly much lower than this (Fig. 4C). This strongly suggests that the fibers have polarity, and grow from only one end, similar to several natural fibers (*8*).

**Fig. 4.**
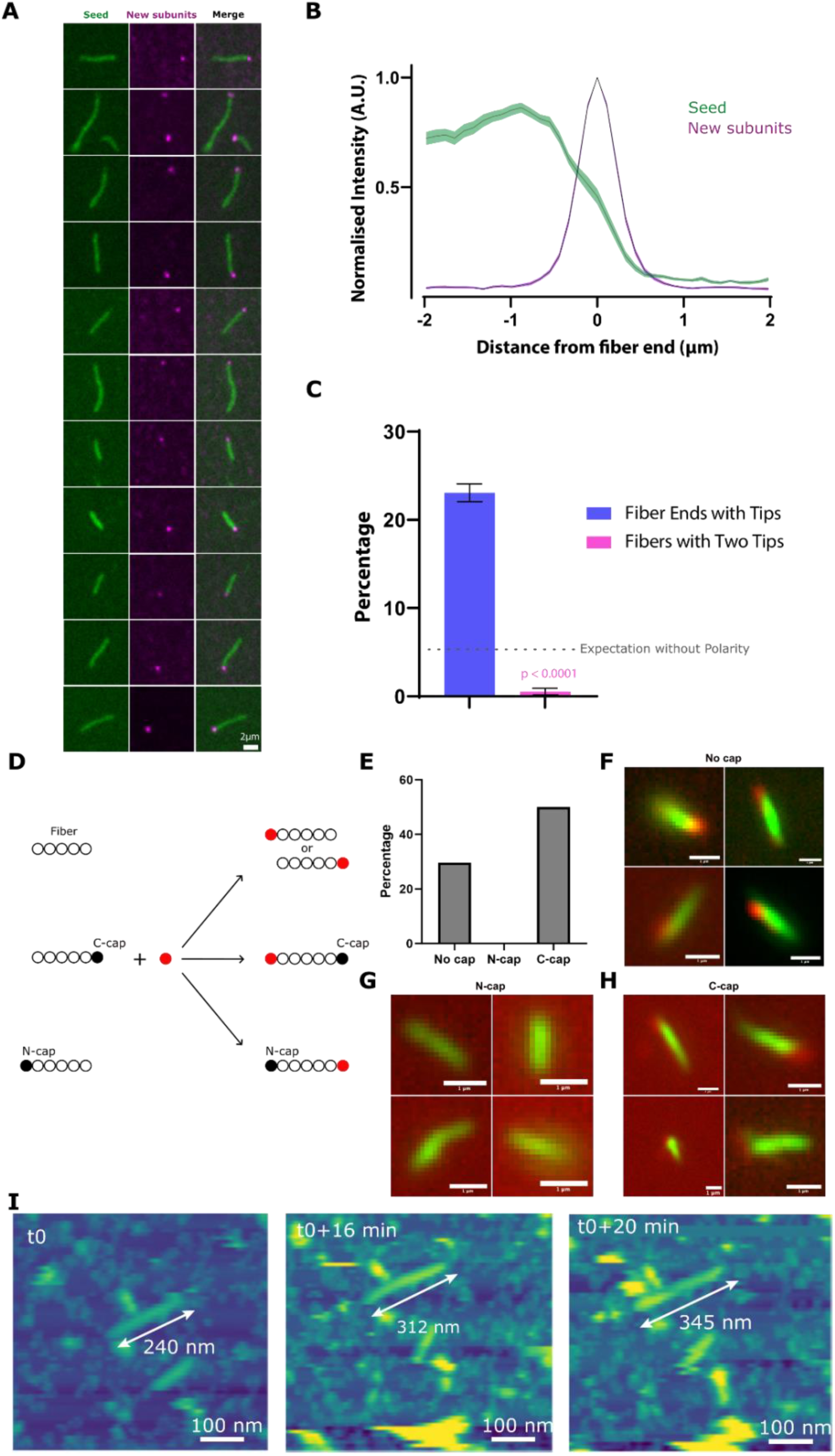
Polarized growth of filaments. (**A)** Fibers were grown using GFP-labeled subunits (HP35 GFP + PLP6-GFP). Additional HP35 subunits and PLP6 peptides labeled with red fluorescent protein mKO2 were then added. The TIRFM images show that the red additions are almost always on just one end of the green fiber seeds. (**B)** Intensity analysis of fiber growth at the end. n=42 fibers. (**C**) Statistical analysis of fiber ends with tips. N = 52 fields of view (FOV), n = 8-21 fibers per FOV. (**D)** Schematic of control experiment of green fiber seed and red new fiber with caps (top) and the design of caps (bottom). (**E**). The percentage of red tip growth quantified from fluorescent images of fiber growth (**F**) without caps and (**G**) with N-cap or (**H**) with C-cap. Scales bars are one micrometer long in **F**-**H**. (**I**)) In-situ AFM images to show the elongation of HP35 protein fiber.

To determine which end the fibers preferentially grow from, we converted the monomeric subunits into fiber capping units by mutating the interfaces that drive fiber formation on one side of the monomer or the other (Fig. S4). We define one of these as the “N-cap” as it contains mutations closer to the monomer N terminus; the other we refer to as the “C-cap”. We first tested the caps by growing new HP35 fiber without seed in the presence of N-cap or C-cap; we found that both inhibit fiber formation (Fig. S5). To determine which end the seeded filaments grow from, we generated green fiber seeds as described above, incubated them with either N-cap, C-cap, or no cap, and then added far-red cy5 labeled new protein and peptide to monitor growth (Fig. 4D). In the absence of cap, 8 red tips were found on 27 fibers; in the presence of N-caps, 0 red tips were found on 19 green fibers, and in the presence of C-cap, 7 red tips were found on 14 fibers, (Fig. 4E-H). We interpret this blocking of fiber extension by the N-caps to indicate that growth from the fiber N-terminal end is strongly favored over growth from the C-terminal fiber end. Atomic Force Microscopy (AFM) images also show that HP35 elongation occurs through the attachment of protein monomers at a single end of the tip, giving rise to directional growth (Fig. 4I).

While protein fibers in biology exhibit nucleation-dependent formation, for example, microtubules are nucleated by the γ-tubulin ring complex, and actin by the Arp2/3 complex, such behavior to our knowledge has not been observed in designed protein fiber systems. A multivalent assembly of monomeric fiber subunits (with a subset of the fiber-forming interactions removed by mutation) was found to capture already existing fibers, but did not promote fiber growth at sub-critical concentrations (*5*). We explored whether assemblies of peptide ligand geometrically matched to the end of the HP35 fiber could nucleate assembly by fusing 4 copies of PLP5 connected by flexible GS linkers to a C3 trimeric oligomer compatible with the fiber structure; we reasoned that this could increase the local concentration of fiber forming-competent monomer and promote nucleation (Fig. 5A). Addition of small amounts of nucleator to HP35 protein and PLP6 peptide substantially increased the rate of fiber formation (Fig. 5B). Green fluorescent labeled nucleator was detectable at the end of the far-red labeled fiber (Fig. 5C). Fiber extension remains dependent on the peptide ligand: addition of nucleator to HP35 alone did not lead to fiber growth (Fig. 5D). AFM characterization also indicated that the nucleator increases both the density and length of HP35 protein fibers without changes in morphology (Fig. 5E, fig S6), with a significant number of fibers observed after 24 hours of incubation with nucleators (Fig. 5F and G). In situ AFM revealed fiber growth from pre-absorbed nucleator (Fig. 5H and I). This dependence of fiber assembly on the nucleator opens up possibilities for control of assembly at specific sites in cells and for amplification of signals in diagnostics assays (a very small amount of nucleator can generate a large assembly that can be read out using fluorescence or other means).

**Fig. 5.**
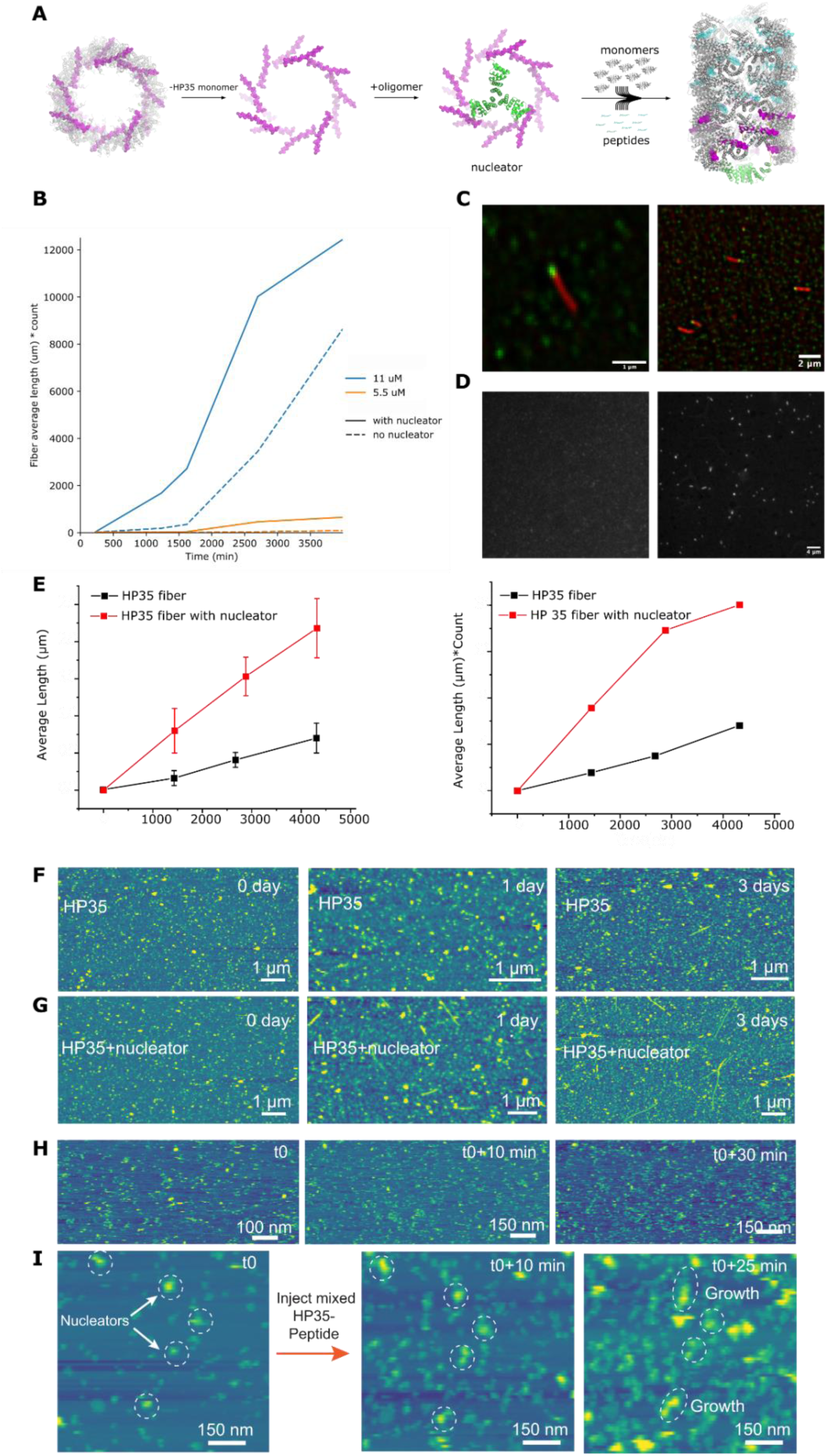
Filament growth with designed nucleator. (**A)** Construction of nucleator with designed oligomers displaying multiple peptides locally. Flexible GS linkers are shown in yellow dashed lines. (**B)** Quantification of fiber growth over time at different concentrations with (solid line) and without (dash line) nucleator from fluorescent microscope images. (**C)** TIRFM images of fibers (red) grown from nucleators (green). (**D**) HP35 mixed with nucleator without (left) and with (right) PLP6 peptide. (**E**) AFM quantification for the average length (left) and total length of HP35 fibers with and without nucleator over time. (**F**) AFM images of HP35 fibers growth without nucleator. (**G**) AFM images of HP35 fibers growth with nucleator. (**H**) In-situ AFM images of HP35 protein without a nucleator over the course of growth time. (**I**) In-situ AFM images of HP35 protein with nucleators. The nucleators were deposited on mica, followed by perfusion with fresh HP35 protein solution.

## Discussion

The assembly of the HP35 fiber is considerably more intricate than previously designed constitutively forming fibers and approaches the complexity of protein filaments of nature. Assembly is allosterically controlled, directional, and sensitive to the addition of nucleators. In this section, we review these properties and discuss their origins.

Assembly of the HP35 fiber is absolutely dependent on the addition of the effector peptide. This control is allosteric, as the peptide is distant from the subunit-subunit interfaces that generate the fiber. This allosteric regulation likely reflects the large hydrophobic groove in the subunit that forms the binding pocket for the effector; in the absence of an effector, this groove may close on itself or drive the formation of transient assemblies, both of which could block fiber assembly. In a manner somewhat similar to the binding of steroid hormones to steroid hormone receptors (*9*), the binding of effector could promote assembly both by driving completion of folding and locking the subunit in a fiber assembly competent state and by disfavoring oligomeric species.

The observation of polarized growth of the HP35 fiber is notable as it requires no external energy input. The energy for adding a new subunit in both directions must be equivalent, and hence the different growing speeds at the two ends must arise from kinetic effects. In contrast to the previously designed filament where the four interfaces are separated (*3*, *5*), one interfacial helix of HP35 contacts the other two adjacent subunits (Fig. 2E) completing an overall triangle-shaped interface. In contrast, the addition of a subunit on the other end leads to the formation of two disconnected half interfaces (Fig. S7). The completion of the triangle interface may be more concerted than the formation of the two half-interfaces, and be kinetically favored. Alternatively, the effector-bound HP35 subunit may have a conformation slightly different from the fiber-integrated form, with a better fit for binding to the N side than the C side.

The nucleation of the HP35 fiber by oligomers presenting large numbers of subunits provides a means to direct fiber formation to specific locations. The nucleator likely functions by increasing the local peptide concentration, creating a large number of peptide-bound HP35 locally that are competent for fiber formation. This genetically encoded nucleator could enable the directing of filament growth to specific locations both in cells and in higher-order materials, and form the basis for signal amplification in detection applications.

More generally, our results suggest that the complex assembly dynamics of many protein filament systems in biology may not have required extensive evolutionary optimization. Our results show that a considerable increase in the richness of assembly behaviors accompanies the transition from constitutive single-component fibers as in our previous work (*5*) to the two-component effector-dependent filament such as the one studied here, likely due to the greater conformational heterogeneity of two-component systems.

## Acknowledgments

We thank S. Gerben, A. Murray, P. Heine, and M. DeWitt for their help with protein production. We thank J Decarreau for help with fluorescent microscopy. We also thank the Arnold and Mabel Beckman Cryo-EM Center at the University of Washington for the use of electron microscopes.

## Author contributions

H.S. and D.B. designed the project. H.S. carried out the design calculations, expressed, purified, and screened design proteins by negative stain EM. E.M.L., H.S. and J.K. carried out the cryo-EM data acquisition and structure determination. C.S. and J.J.D.Y. carried out the AFM data acquisition and analysis. J.L.W. and E.D. carried out TIRF fluorescence microscopy on the directional growth experiment. H.S. performed fluorescence microscopy for fiber in solution and quantification. H.B. and H.S. designed the polar version of fiber. K.W. and H.S. designed the nucleator. Y.L., H.S., and C.J.K. construct and express protein in mammalian cells. H.S., W.S., and D.B. designed the method figure. H.S. and D.B. wrote the manuscript.

## Methods

### Computational Design Strategy

The helical docking and design method (*5*) was applied to RPB_PLP3_R6 to generate models of helical filament designs. Design trajectories were filtered using the following criteria: a Rosetta Energy Unit difference greater than -15.0 between the polymeric (bound) and monomeric (unbound) states, an interface surface area over 700 Å², Rosetta shape complementarity above 0.62, and fewer than 5 unsatisfied polar residues. Designs meeting these criteria underwent manual refinement, with non-contributory mutations reverted to their original state. The top-scoring design for each configuration was then included in a final protein set for experimental validation.

To tune and facilitate the exchangeability of peptides on the original HP35 fiber, we designed and screened a series of HP35 polar variants, in the presence of shorter PLP repeating peptides, especially (PLP)3 and (PLP)4. In brief, the asymmetry unit of HP35-(PLP)6 fiber model was extracted and served as the input for redesign. The PLP repeating peptide was shortened from six repeats to three or four repeats. The newly exposed surface residues along the peptide binding interface on the HP35 monomer were identified and redesigned by Rosetta FastDesign, in the context of the original fiber filament symmetry. The design process was guided by the Rosetta score function “REF2015”. For the successful design HP35_B3, the interface surface residues “36, 37, 40, 41, 44, 84, 87, 88, 91, 200, 247, 248, 251, 292, 299” were chosen for redesign.

### Protein Expression and Purification

Synthetic genes for 69 designs were optimized for expression in *Escherichia coli* and synthesized by IDT. These genes were cloned into the pET29b+ based vectors with either N-or C-terminal GFP fusion, inserted between the NdeI and XhoI restriction sites. The constructs were transformed into BL21* (DE3) *E. coli* competent cells and grown in 50 ml of Terrific Broth medium containing 200 mg L^-1^ kanamycin. Protein expression, driven by a T7 promoter, proceeded for 24 hours at 37°C using Studier autoinduction(*10*), after which cultures were harvested by centrifugation. Cell pellets were resuspended in Tris-buffered saline (TBS) and lysed with Bugbuster detergent. The soluble fraction, clarified by centrifugation, was purified using Ni^2+^-immobilized metal affinity chromatography (IMAC) on Ni-NTA Superflow resin. The resin-bound lysate was washed with 10 column volumes of 40 mM imidazole and 500 mM NaCl, then eluted with 400 mM imidazole and 75 mM NaCl. Both soluble and insoluble fractions were analyzed by SDS-PAGE. Samples showing protein bands at the expected molecular weight were selected for electron microscopy screening. For further characterization, selected designs were scaled up to 0.5 L, expressed under the same conditions for 24 hours at 37°C, and harvested by centrifugation. Cell pellets were resuspended in TBS, lysed via microfluidization, and purified using the same method.

### Gene Construction for Designed Oligomeric Peptides Nucleator

The protein sequences were engineered for optimal expression in E. coli. Linear DNA fragments encoding these designs were synthesized (eBlocks, Integrated DNA Technologies) with overhangs designed for compatibility with Golden Gate cloning into the LM671 expression vector. This vector contains a Kanamycin resistance gene, a ccdB selection marker flanked by BsaI restriction sites, and a C-terminal superfold-GFP fusion followed by the C-terminal hexahistidine (6xHis) tag for purification, separated by either a PAS or GGSGSG linker.

### Protein Production and Purification (Small Scale)

Linear DNA sequences encoding the designs were introduced into the LM671 vector via Golden Gate assembly, performed in 96-well PCR plates with a 4 µL reaction volume. One microliter of the resulting reaction was then used to transform chemically competent E. coli BL21 (DE3) cells. After a one-hour recovery period in 100 µL of SOC medium, the transformation mixtures were transferred to 96-deep well plates containing 900 µL of LB media with Kanamycin. Cultures were incubated overnight at 37°C. To induce protein expression, 100 µL of each overnight culture was added to fresh 96-deep well plates containing 900 µL of auto-induction media (TBII media with Kanamycin, 2 mM MgSO₄, and 1X 5052) and incubated at room temperature overnight.

Cells were collected by centrifugation at 4000 x g for 15 minutes, and bacterial pellets were lysed in 100 µL of lysis buffer (1X BugBuster (Millipore #70921-4), 0.01 mg/mL DNase, and 1 Pierce protease inhibitor tablet per 50 mL) while shaking at 220 RPM for 30 minutes. After lysates were clarified by centrifugation at 4000 x g for 10 minutes, proteins were purified using Ni-charged MagBeads (GenScript #L00295). The wash buffer consisted of 25 mM Tris pH 8.0, 300 mM NaCl, and 10 mM Imidazole, while the elution buffer contained 25 mM Tris pH 8.0, 300 mM NaCl, and 500 mM Imidazole. Proteins were washed twice with 250 µL of wash buffer and eluted with 120 µL of elution buffer.

### Negative Stain Electron Microscopy

The soluble fractions were concentrated in a buffer (25 mM Tris, 75 mM NaCl, pH 8) for electron microscopy screening. A 6 µL droplet (1 µL sample immediately diluted with 5 µL buffer) was placed onto negatively glow-discharged, carbon-coated 200-mesh copper grids, washed with Milli-Q water, and stained with 0.75% uranyl formate (pH 4.0) as previously described (*11*). The screening was performed using a 120 kV Talos L120C transmission electron microscope (ThermoFisher). Images were captured using a bottom-mounted Teitz CMOS 4k camera system and processed for enhanced contrast with Fiji software(*12*)

### Cryo-Electron Microscopy

CryoEM samples were prepared by applying protein to CFLAT 2/2 holey-carbon grids, blotting away liquid, and plunging into liquid ethane using a Vitrobot (ThermoFisher). Movies were acquired on a Glacios microscope (ThermoFisher) equipped with a K-2 Summit Direct Detect camera (Gatan Inc.) operating in counting mode, with a pixel size of 1.16 Å/pixel, 50 frames, and a total dose of 65 electrons/Å^2^. Automated data collection was performed using Leginon (*13*). The data processing workflow is summarized in Fig. S8. Movies were aligned using the Relion (*7*) implementation of MotionCor2 (*14*), and CTF parameters were estimated using Gctf (*15*). Helices were picked manually from a subset of images and subjected to 2D classification to generate templates for subsequent automated picking in Relion. Particles from automated picking were subjected to further 2D classification using Relion. *Ab initio* reconstruction in cisTEM (*6*), using a subset of particles from automated picking, was used to generate an initial 3D map from which initial helical symmetry parameters were also estimated. Helical auto-refinement was then performed in Relion using the *ab initio* map as a starting reference, with local searches of helical rise and rotation parameters enabled. Bayesian polishing as well as beamtilt, anisotropic magnification, and defocus refinement were performed in Relion. Density modification was performed using ResolveCryoEM in Phenix (*16*, *17*). Atomic models were refined into cryoEM maps using ISOLDE(*18*), followed by real-space refinement in Phenix, with rotamer and Ramachandran restraints disabled and with reference restraints imposed by the input starting model. Cryo-EM data collection, refinement, and validation statistics are summarized in Table S1.

### Fluorescent Microscopy

#### Sample preparation

1uL of fiber solution was diluted in 1mL of TBS buffer (25 mM Tris, 75 mM NaCl, pH 8), and 50 uL of diluted sample was added to the well for imaging.

#### Image acquisition

##### IN Cell Analyzer 2500HS

Fibers were attached directly to the glass surface of a 384-well plate (Greiner Bio-one, cat# 781892, https://www.gbo.com/) and allowed to stick for 5 minutes before washing to remove the background signal. Images were automatically sampled using an IN Cell Analyzer 2500HS and a 60X 0.95 NA Plan Apo CFI objective (Nikon, 12605). Fibers were illuminated with a seven-color Solid State Illuminator (SSI) for fluorescence excitation. Fluorescent signals were acquired sequentially using the following filter setup: Green (excitation 473/28 nm, emission 511/23 nm), Red (excitation 57530 nm, emission 623/42 nm), and Far-red (excitation 631/28 nm, emission 684/24 nm). Imaging was controlled using the IN Cell Analyzer 2500 HS software version 7.4 and the light was collected on a sCMOS camera without binning. Between 15-25 fields of view spaced evenly throughout the well were imaged per condition.

##### OMX

Fibers were attached directly to the glass surface of either an Ibidi µ-Slide 0.5 or an Ibidi 18-well glass-bottom coverslide (cat# 806076, 81817, respectively. Ibidi, https://ibidi.com/). Two-color images of fibers were acquired with a commercial OMX-SR system (Leica-microsystems) equipped with an Olympus Apo N 60× 1.49 NA TIRF oil immersion lens. Toptica diode lasers with wavelengths of 405nm, 488nm, and 640nm were used for excitation. Emission was collected on three separate PCO.edge sCMOS cameras. 256×256 images (pixel size 6.5 μm) were captured with no binning. The acquisition was controlled with AcquireSR Acquisition control software. Images from different color channels were registered in SoftWoRx using parameters generated from a gold grid registration slide (GE Healthcare).

##### Image analysis

To segment the fiber images, a subset of 35 images were set aside for training a random forest classifier in a custom Jupyter notebook. The training images were manually annotated to define true positive fiber pixels in each image. The manual annotations were then used to train a classifier using the RandomForrestClassifier module from the Scikit-learn library (*19*). Our model defined the max tree depth as 10 and the number of estimators to be 50. Accuracy scores > 0.99 were measured in our training set and the results of this custom model were qualitatively similar to a previously generated model using the Weka trainable segmentation module in Fiji (*12*, *20*). Following model training a second analysis notebook was used to segment each image. After model-generated segmentation, we processed the output by removing small objects (less than 64 pixels) and performed a binary closing operation to prevent over-segmentation. The resulting objects were measured for size and intensity.

### Cell culture

To enable controllable expression of the multiple fiber components, we put gene sequences encoding the HP35-GFP and the BFP-FNC19 nucleator under the control of a cumate-regulated promoter (cloned into the pCDH-CuO-MCS vector, System Biosciences, #QM500A-1) and a tetracycline-response element (TRE-HP35-GFP-Ubc-rtTA), respectively, while the gene encoding PLP6-mScarI was cloned into the pcDNA3.1(+) vector.

HeLa and HEK293T cells were cultured in DMEM (Gibco) supplemented with 10% HyClone FetalClone II (FC-II, Cytiva SH30066.03) and 1% penicillin-streptomycin (Gibco) at 37 °C with 5% CO2. To enable stable expression of the cumate-responsive switch for inducible expression of the fiber, lentivirus was prepared from the lentiviral expression vector encoding the CymR gene (System Biosciences, #QM200PA-2) using HEK293T cells. The HEK293T cells were co-transfected with the expression vector and the packaging plasmids (psPAX2 and pMD2.G) using Lipofectamine 3000 following manufacturer’s instructions. The supernatant containing the viral particles was harvested 72 hours after transfection and filtered through 0.45 µm surfactant-free cellulose acetate (SFCA) filters (Corning #431220). HeLa cells were transduced with the collected viral supernatant in the presence of 8 µg/mL polybrene (Santa Crutz Biotechnology). The transduced cells were then subjected to puromycin selection for a week.

Thereafter, the cells constitutively expressing CymR were seeded into glass-bottom 96-well cell culture plates (Cellvis P96-1.5H-N) at a density of ∼40,000 cells per well. The following day, cells were transfected using Lipofectamine 3000 according to the manufacturer’s instructions with all three plasmids (HP35 protein, PLP6 peptide and nucleator) mixed at a molar ratio of 2:2:5 and 2:2:1. The medium was exchanged to the DMEM culture medium without doxycycline (we found that a minimal expression level of the FNC19 nucleator in the absence of doxycycline was sufficient to induce fiber formation), and that with varying concentrations of cumate (2, 10 or 50 µg/mL) 24 and 48 hours after transfection, respectively. Cells were imaged 72 hours after transfection in HEPES-buffered solution (10 mM HEPES, 140 mM NaCl, 5 mM KCl, 2 mM MgCl2, 2 mM CaCl2, 10 mM glucose, 15 mM sucrose, pH 7.4 adjusted by NaOH) on InCell Analyzer. (Images were imported into Fiji and processed.)

### Directional growth experiment

HP35-GFP (30uM) and PLP6-GFP (30uM) were incubated at room temperature for 48 hours in 25mM Tris, 75mM NaCl, 1mM TCEP. After 48 hours, HP35-cys (30uM) and PLP6-mKO2 (30uM) were separately mixed in 25mM Tris, 75mM NaCl, 1mM TCEP. The pre-assembled HP35-GFP/PLP6-GFP fibres were added at a 1:5 v/v ratio. This mixture was incubated for 1 hour, before 1:20 dilution with 25mM Tris, 75mM NaCl, 1mM TCEP, 0.4% methlycellulose. 20ul was added to an Ibidi flow cell mounted on clean room-grade coverslips (custom, 25 × 75 mm2, Nexterion), and passivated with PLL-PEG (0.1 mg ml−1 in 20 mM Hepes, pH 7.6; 5 min). Fibers were allowed to deposit on the glass for 15 minutes, before washing with and TIRF imaging in 25mM Tris, 75mM NaCl, 1mM TCEP, 0.4% methlycellulose.

### Atomic Force Microscopy

The HP35 protein solution was diluted to 1 μM using a KCl-Tris buffer (pH 7.4; KCl: 500 mM, Tris: 21 mM). A 70 μL aliquot of the protein solution was incubated on a mica surface, which was then placed into the AFM liquid cell. For the perfusion AFM mode, maintain a perfusion rate of 0.1 mL/min. All AFM images were acquired in tapping mode using a Cypher ES AFM (Asylum Research). Carbon/Diamond-like Carbon probes (USC-F1.2-k0.15) from NanoWorld were employed for all imaging. AFM height and phase images of SF were captured at a scan frequency of 1.5 Hz. Height images were processed using Gwyddion SPM data analysis software.

---

**Fig. S1.**
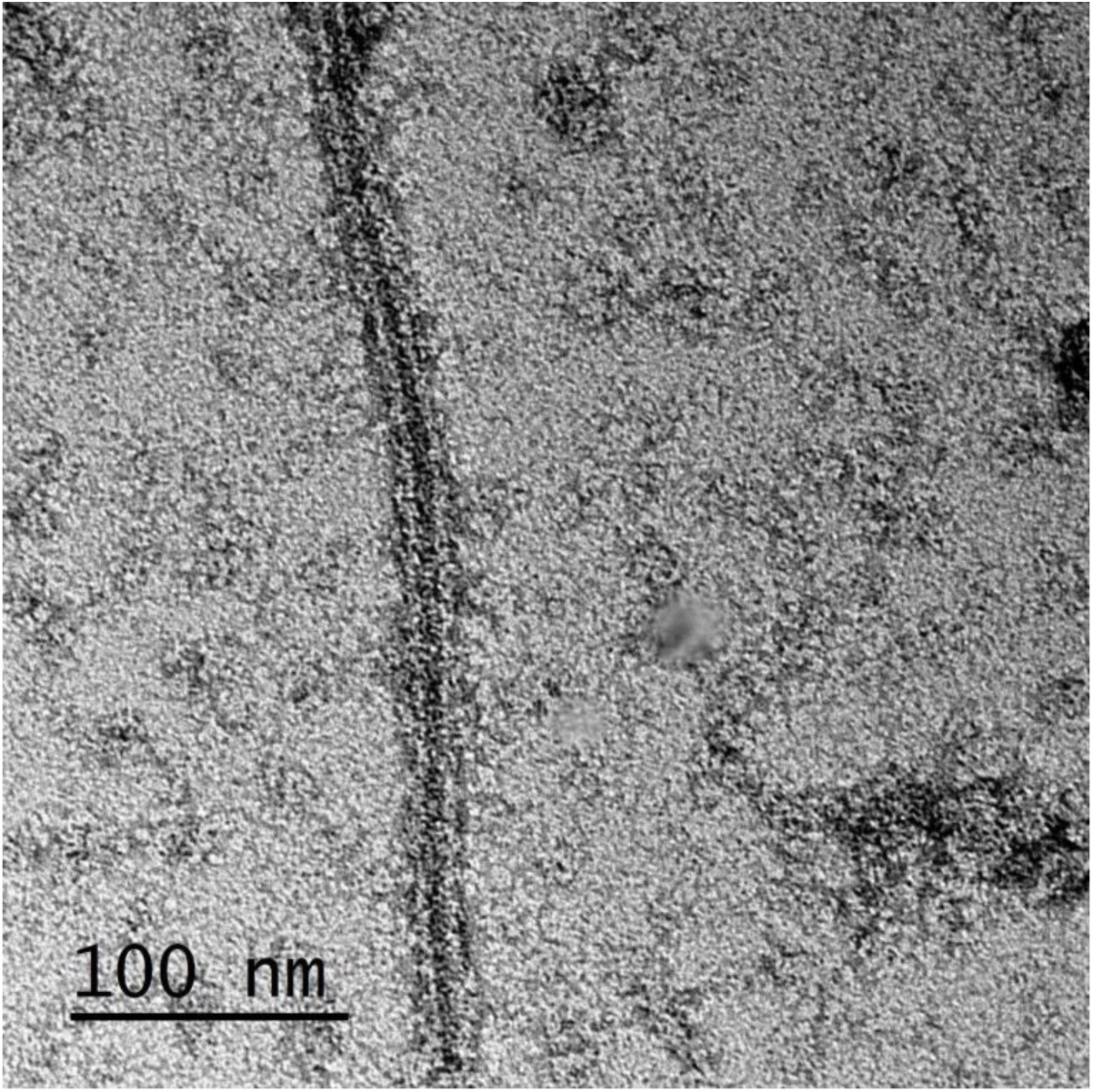
Negative stain EM image of HP35 fiber

**Fig. S2.**
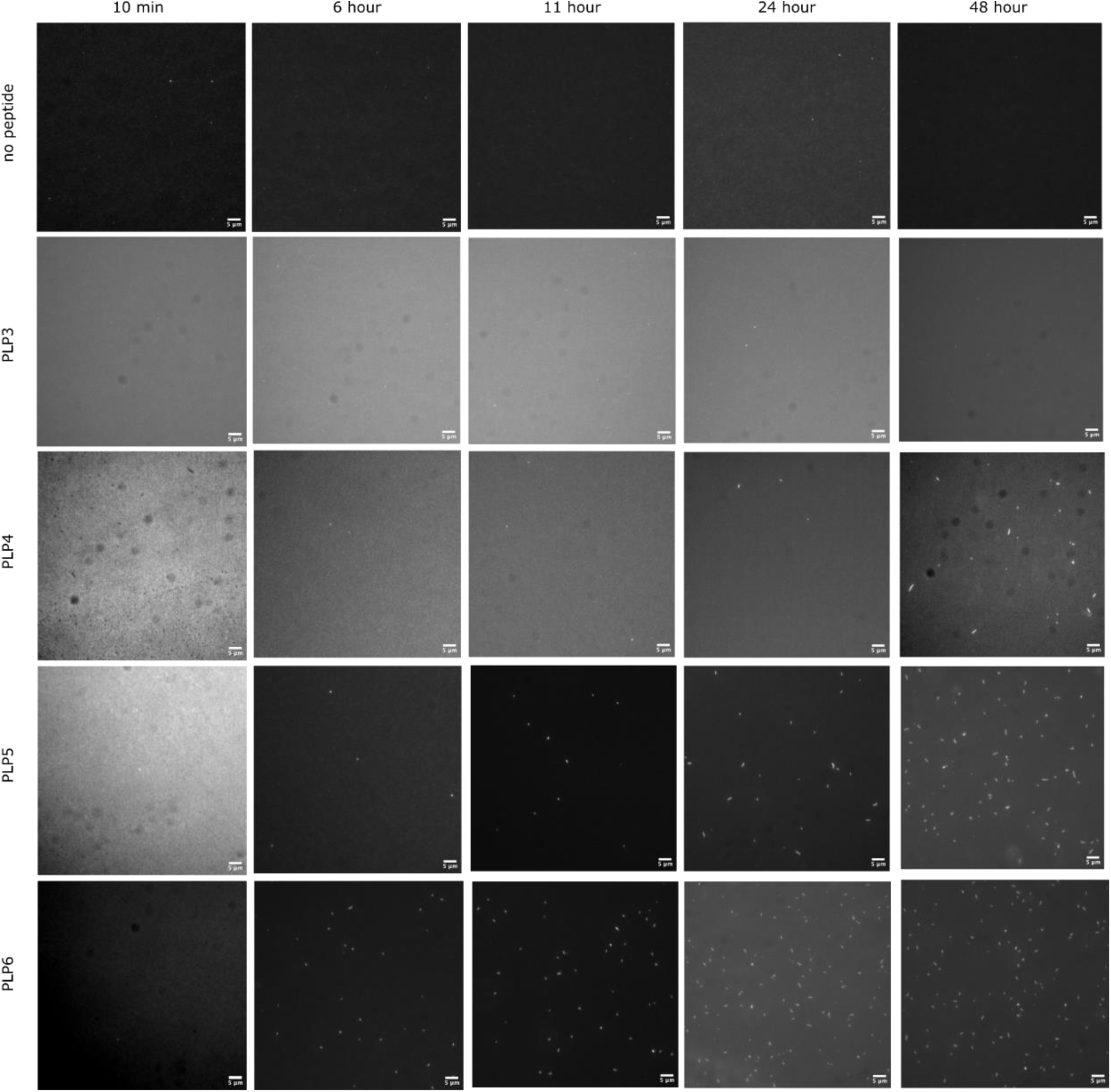
Filament growth dynamic. Representative fluorescent microscopy images of HP35 mixed with different lengths of peptide PLP3-5 over time

**Fig. S3.**
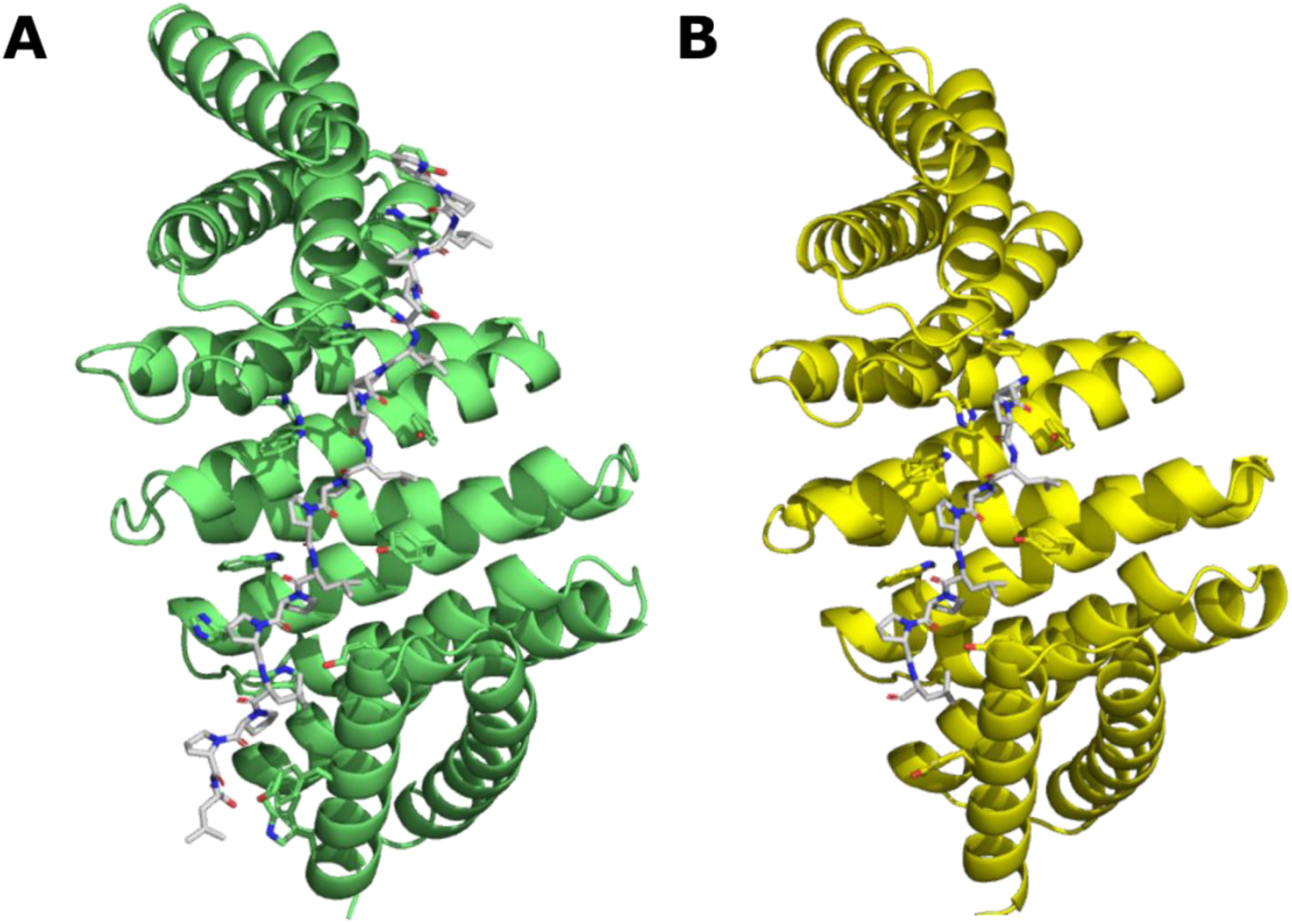
HP35_B3 design model. (**A**) Design model of HP35 and (**B**) Design model of HP35_B3

**Fig. S4.**
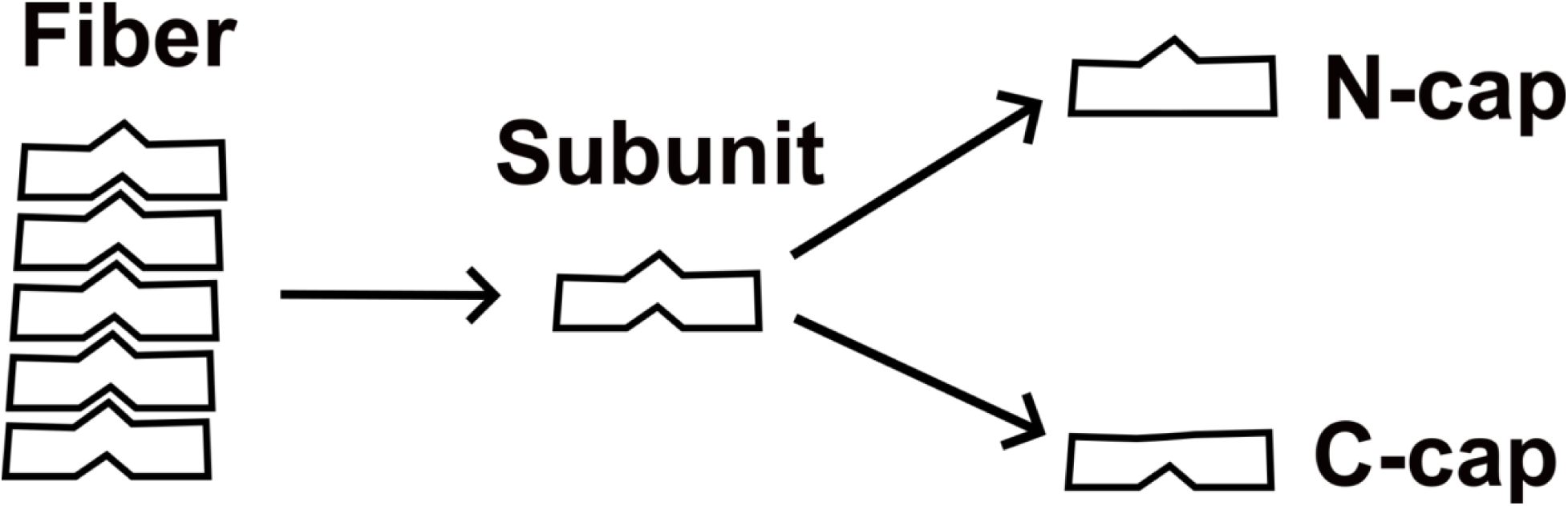
Fiber cap design. Design of N-cap and C-cap by knocking out different halves of the design interfaces

**Fig. S5.**
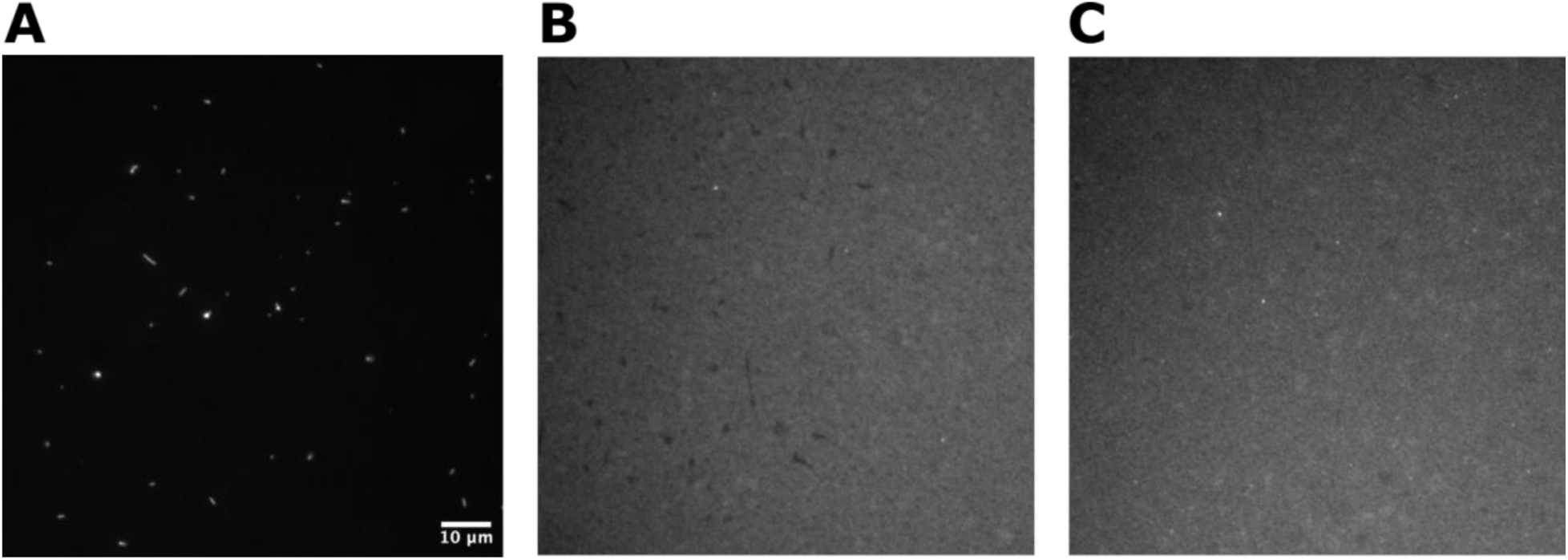
Fiber growth in the presence of caps. HP35 mixed with PLP6 peptide in the absence of caps **(A)** and the presence of N-cap (**B**) or C-cap (**C**).

**Fig. S6.**
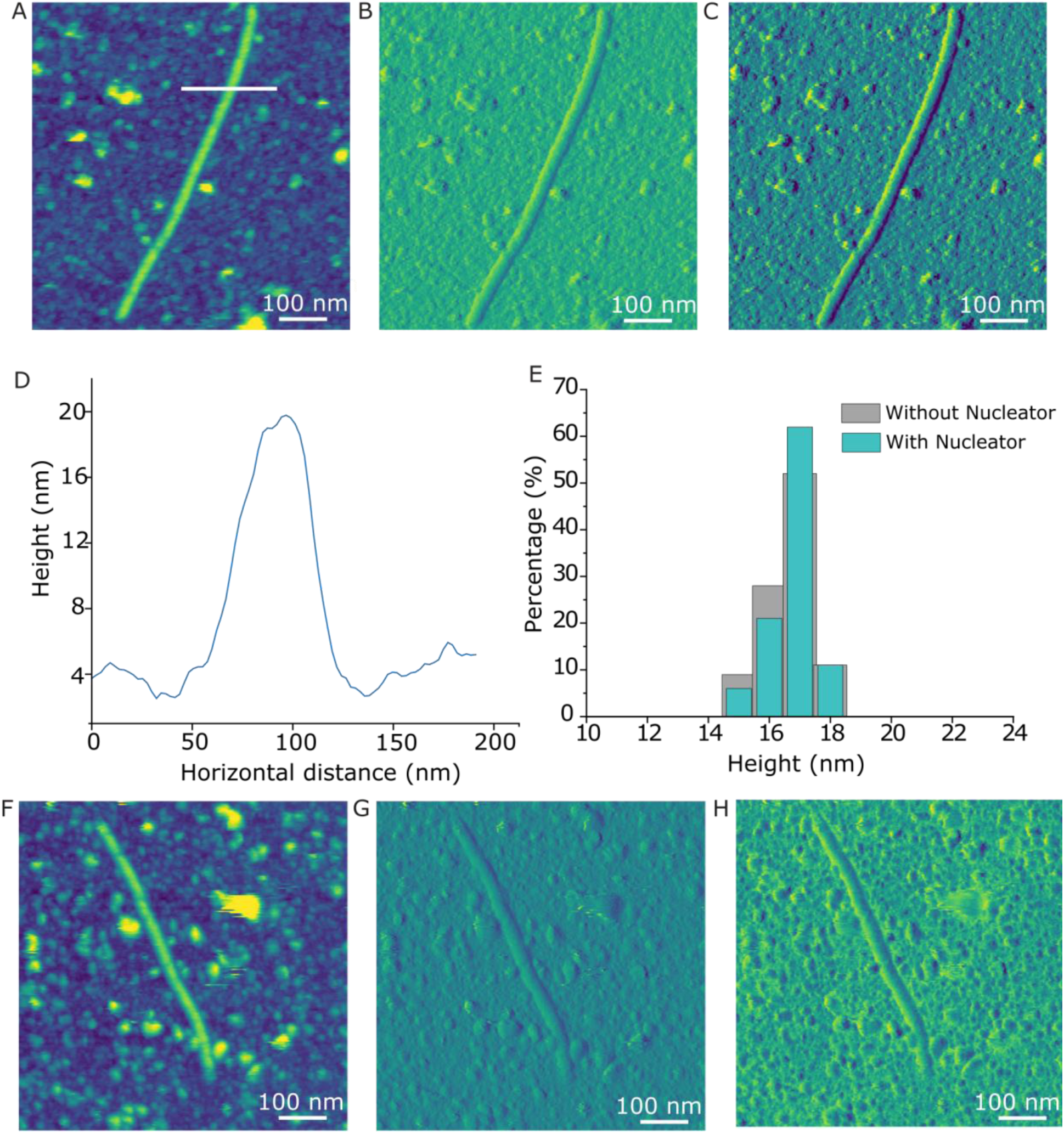
Morphology of HP35 Protein Fibers. (**A-C**) Topography (**A**), amplitude (**B**), and phase (**C**) images of HP35 fibers with nucleator. (**D**) Height profile along the white line in image A. (**E**) Height distribution of the protein fibers. (**F-H**) Topography (**F**), amplitude (**G**), and phase (**H**) images of HP35 fibers without nucleator. The HP35 protein fibers exhibit no significant differences in their final morphology with or without nucleator (A-C, F-H), with both samples displaying a smooth surface and straight contour. The height of the fibers is centered around 17 nm (D-E), which aligns with the designed model.

**Fig. S7.**
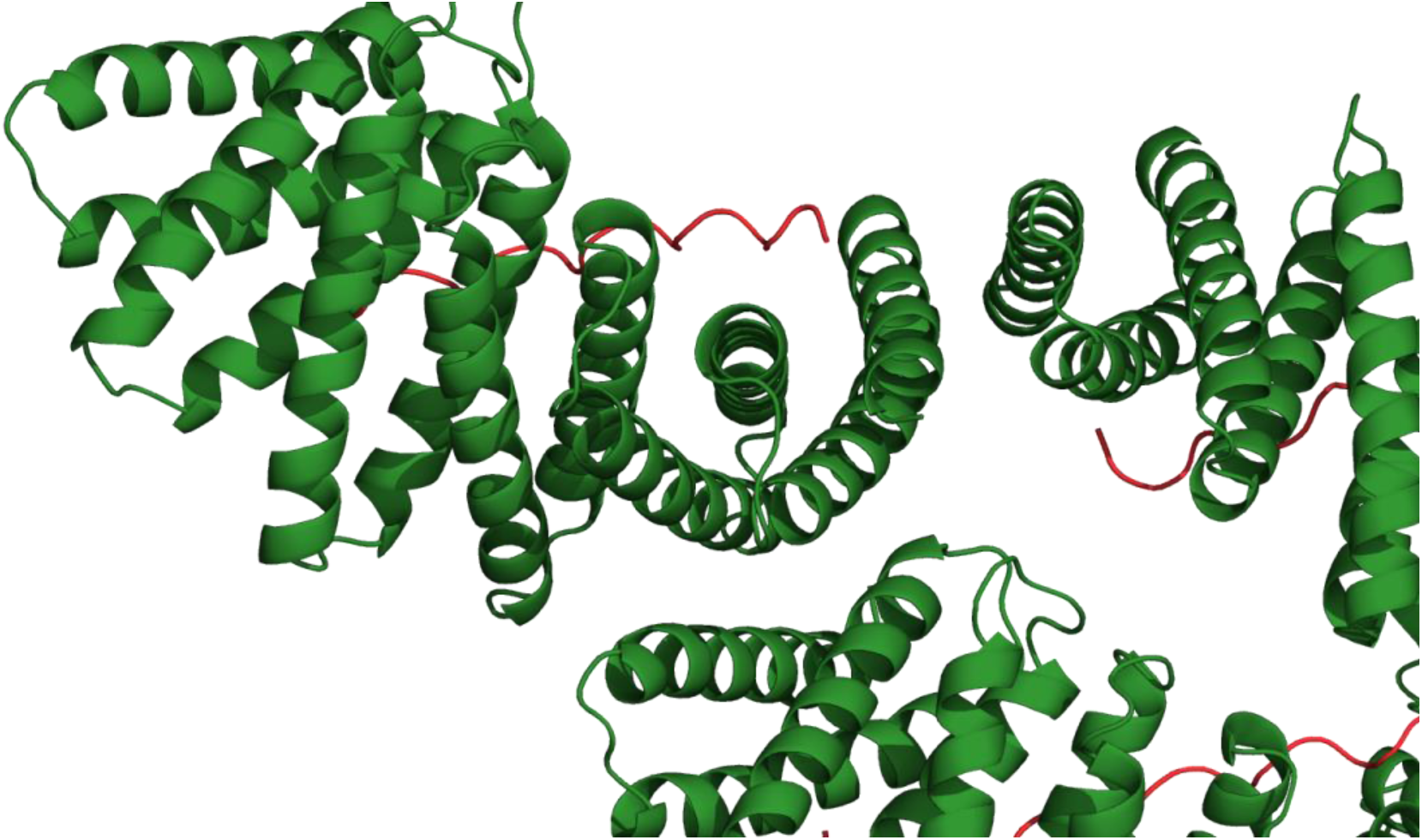
Formation of two disconnected half interfaces with the addition of a subunit on the other end

**Fig. S8.**
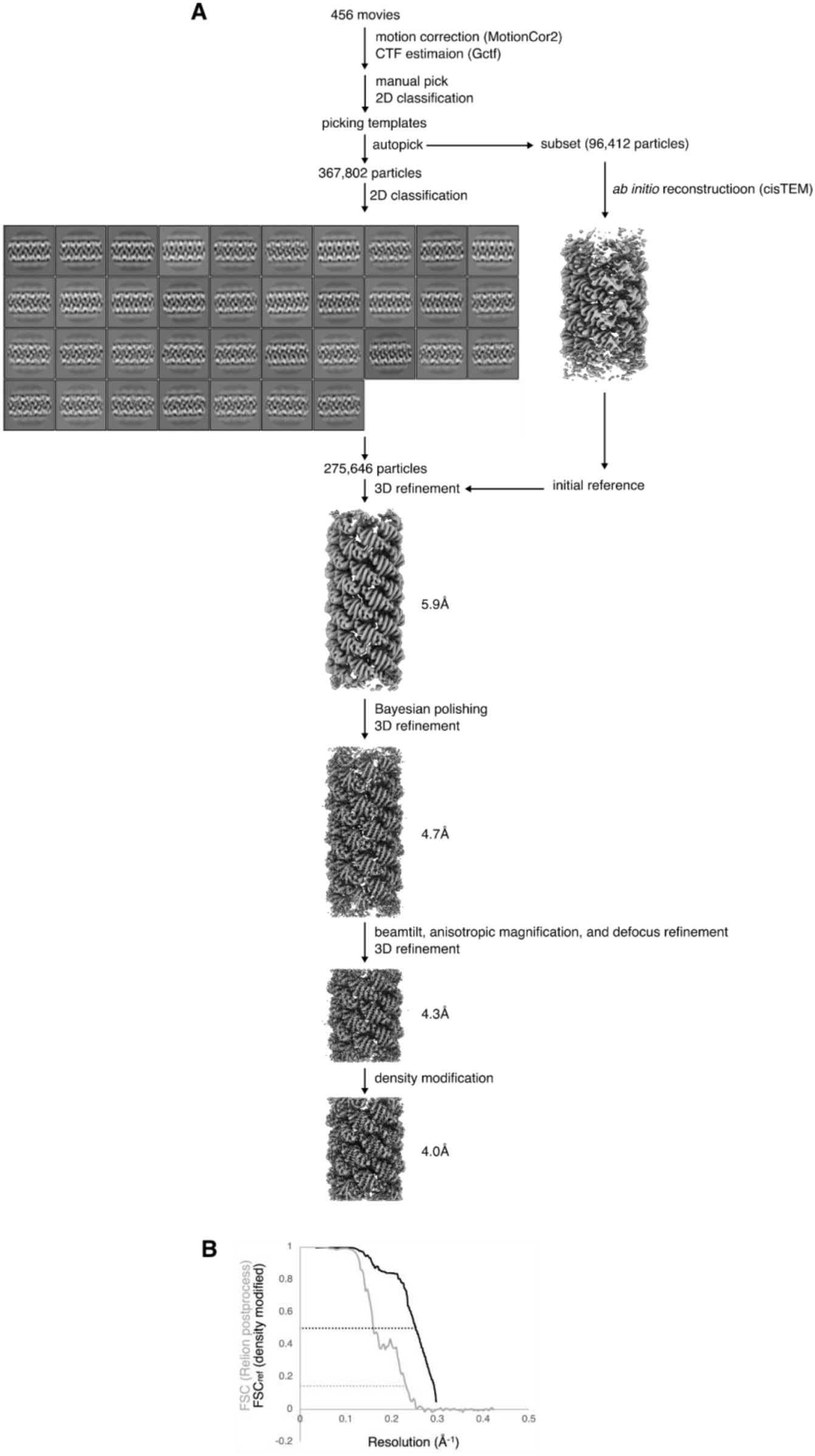
CryoEM data processing. (A) Flowchart of cryoEM data processing. (B) Half-map FSC curve from Relion postprocessing (grey) and FSCref curve after density modification (black) and corresponding resolution estimates.

**Table S1:** Cryo-EM data collection, refinement and validation statistics

|  |  |
| --- | --- |
| Structure | HP35 filament |
| PDB code | 8VIM |
| EMDB code | 43266 |
| Magnification | 36000 |
| Voltage (kV) | 200 |
| Electron fluence (e-/Å<sup>2</sup>) | 65 |
| Defocus range (µm) | 0.6-2.2 |
| Pixel size (data collection) (Å) | 1.16 |
| Pixel size (reconstruction) (Å) | 1.16 |
| Point group symmetry | C1 |
| Helical rise (Å) | 6.2 |
| Helical rotation (degrees) | 132 |
| Particle images (no.) | 275,646 |
| Resolution (0.143 FSC) (Å) | 4.3 |
| Resolution, density modified (0.5 FSC<sub>ref</sub>) (Å) | 4 |
| R.m.s. deviation bond lengths (Å) | 0.42 |
| R.m.s. deviation bond angles (°) | 0.66 |
| MolProbity score | 1.16 |
| Clashscore | 2 |
| C-beta deviations | 0 |
| Rotamer outliers (%) | 0 |
| Ramachandran favored (%) | 97 |
| Ramachandran allowed (%) | 2 |
| Ramachandran outliers (%) | 0 |

## References

1. E. Nogales, H.-W. Wang, Structural intermediates in microtubule assembly and disassembly: how and why? Curr. Opin. Cell Biol. 18, 179–184 (2006).

2. T. D. Pollard, G. G. Borisy, Cellular motility driven by assembly and disassembly of actin filaments. Cell 112, 453–465 (2003).

3. H. Shen, E. M. Lynch, S. Akkineni, J. L. Watson, J. Decarreau, N. P. Bethel, I. Benna, W. Sheffler, D. Farrell, F. DiMaio, E. Derivery, J. J. De Yoreo, J. Kollman, D. Baker, De novo design of pH-responsive self-assembling helical protein filaments. Nat. Nanotechnol., doi: 10.1038/s41565-024-01641-1 (2024).

4. K. Wu, H. Bai, Y.-T. Chang, R. Redler, K. E. McNally, W. Sheffler, T. J. Brunette, D. R. Hicks, T. E. Morgan, T. J. Stevens, A. Broerman, I. Goreshnik, M. DeWitt, C. M. Chow, Y. Shen, L. Stewart, E. Derivery, D. A. Silva, G. Bhabha, D. C. Ekiert, D. Baker, De novo design of modular peptide-binding proteins by superhelical matching. Nature, doi: 10.1038/s41586-023-05909-9 (2023).

5. H. Shen, J. A. Fallas, E. Lynch, W. Sheffler, B. Parry, N. Jannetty, J. Decarreau, M. Wagenbach, J. J. Vicente, J. Chen, L. Wang, Q. Dowling, G. Oberdorfer, L. Stewart, L. Wordeman, J. De Yoreo, C. Jacobs-Wagner, J. Kollman, D. Baker, De novo design of self-assembling helical protein filaments. Science 362, 705–709 (2018).

6. T. Grant, A. Rohou, N. Grigorieff, cisTEM, user-friendly software for single-particle image processing. Elife 7 (2018).

7. S. H. W. Scheres, RELION: implementation of a Bayesian approach to cryo-EM structure determination. J. Struct. Biol. 180, 519–530 (2012).

8. P. J. Carman, K. R. Barrie, G. Rebowski, R. Dominguez, Structures of the free and capped ends of the actin filament. Science 380, 1287–1292 (2023).

9. G. F. Allan, X. Leng, S. Y. Tsai, N. L. Weigel, D. P. Edwards, M. J. Tsai, B. W. O’Malley, Hormone and antihormone induce distinct conformational changes which are central to steroid receptor activation. J. Biol. Chem. 267, 19513–19520 (1992).

10. F. W. Studier, Protein production by auto-induction in high density shaking cultures. Protein Expr. Purif. 41, 207–234 (2005).

11. B. L. Nannenga, M. G. Iadanza, B. S. Vollmar, T. Gonen, Overview of Electron Crystallography of Membrane Proteins: Crystallization and Screening Strategies Using Negative Stain Electron Microscopy. Curr. Protoc. Protein Sci. 72, 17.15.1–17.15.11 (2013).

12. J. Schindelin, I. Arganda-Carreras, E. Frise, V. Kaynig, M. Longair, T. Pietzsch, S. Preibisch, C. Rueden, S. Saalfeld, B. Schmid, J.-Y. Tinevez, D. J. White, V. Hartenstein, K. Eliceiri, P. Tomancak, A. Cardona, Fiji: an open-source platform for biological-image analysis. Nat. Methods 9, 676–682 (2012).

13. C. Suloway, J. Pulokas, D. Fellmann, A. Cheng, F. Guerra, J. Quispe, S. Stagg, C. S. Potter, B. Carragher, Automated molecular microscopy: the new Leginon system. J. Struct. Biol. 151, 41–60 (2005).

14. S. Q. Zheng, E. Palovcak, J.-P. Armache, K. A. Verba, Y. Cheng, D. A. Agard, MotionCor2: anisotropic correction of beam-induced motion for improved cryo-electron microscopy. Nat. Methods 14, 331–332 (2017).

15. K. Zhang, Gctf: Real-time CTF determination and correction. J. Struct. Biol. 193, 1–12 (2016).

16. T. C. Terwilliger, S. J. Ludtke, R. J. Read, P. D. Adams, P. V. Afonine, Improvement of cryo-EM maps by density modification. Nat. Methods 17, 923–927 (2020).

17. P. D. Adams, P. V. Afonine, G. Bunkóczi, V. B. Chen, I. W. Davis, N. Echols, J. J. Headd, L.-W. Hung, G. J. Kapral, R. W. Grosse-Kunstleve, A. J. McCoy, N. W. Moriarty, R. Oeffner, R. J. Read, D. C. Richardson, J. S. Richardson, T. C. Terwilliger, P. H. Zwart, PHENIX: a comprehensive Python-based system for macromolecular structure solution. Acta Crystallogr. D Biol. Crystallogr. 66, 213–221 (2010).

18. T. I. Croll, ISOLDE: a physically realistic environment for model building into low-resolution electron-density maps. Acta Crystallogr D Struct Biol 74, 519–530 (2018).

19. F. Pedregosa, G. Varoquaux, A. Gramfort, V. Michel, B. Thirion, O. Grisel, M. Blondel, G. Louppe, P. Prettenhofer, R. Weiss, R. J. Weiss, J. Vanderplas, A. Passos, D. Cournapeau, M. Brucher, M. Perrot, E. Duchesnay, Scikit-learn: Machine Learning in Python. J. Mach. Learn. Res. **abs**/1201.0490 (2011).

20. I. Arganda-Carreras, V. Kaynig, C. Rueden, K. W. Eliceiri, J. Schindelin, A. Cardona, H. Sebastian Seung, Trainable Weka Segmentation: a machine learning tool for microscopy pixel classification. Bioinformatics 33, 2424–2426 (2017).

